# High sensitivity (zeptomole) detection of BODIPY heparan sulfate (HS) disaccharides by ion-paired RP-HPLC and LIF detection enables analysis of HS from mosquito midguts

**DOI:** 10.1101/2020.01.21.913954

**Authors:** Marissa L. Maciej-Hulme, Anaëlle C. N. Leprince, Andre Lavin, Scott E. Guimond, Jeremy E. Turnbull, Julien Pelletier, Edwin A. Yates, Andrew K. Powell, M. A. Skidmore

## Abstract

The fine structure of heparan sulfate (HS), the glycosaminoglycan polysaccharide component of cell surface and extracellular matrix HS proteoglycans, coordinates the complex cell signalling processes that control homeostasis and drive development in multicellular animals. In addition, HS is involved in the infection of mammals by viruses, bacteria and parasites. The current detection limit for fluorescently labelled HS disaccharides that is in the low femtomole range (10^-15^ mol), has effectively hampered investigations of HS composition from small, functionally-relevant populations of cells and tissues. Here, an ultra-high sensitivity method is described that utilises a combination of reverse-phase HPLC, with tetraoctylammonium bromide (TOAB) as the ion-pairing reagent and laser-induced fluorescence detection of BODIPY-FI-labelled disaccharides. The method provides an unparalleled increase in the sensitivity of detection by ∼ six orders of magnitude, to the zeptomolar range (∼10^-21^ moles), enabling detection of <1000 labelled molecules. This facilitates determination of HS disaccharide compositional analysis from minute biological samples, as demonstrated by analysis of HS isolated from the midguts of *Anopheles gambiae* mosquitoes that was achieved without approaching the limit of detection.

## Introduction

Heparan sulfate (HS) is a linear, anionic glycosaminoglycan (GAG) polysaccharide component of cell surface and extracellular matrix HS proteoglycans (HSPGs), whose fine structure dictates coordination of the complex cell signalling processes that control homeostasis and drive development in multicellular animals. HS, which is displayed at the mammalian cell surface, is also known to interact with viruses (e.g. HIV ^1^ and Zika virus ^2, 3^) and other cells, including pathogenic microorganisms (e.g. *Toxoplasma gondii* ^4, 5^, *Plasmodium falciparum* ^5, 6^, and *Leishmania* parasites ^7–9^) and is often involved in the process of infection. In addition, diffusible HS oligosaccharide fragments released by heparanase activity are thought to exert influence further afield ^10^.

The biosynthesis of HS occurs in the endoplasmic reticulum and Golgi, where the nascent chain is modified during *de novo* synthesis on the protein core. Specific enzymes either transfer sulfate groups (N-deacetylase/sulfotransferases, 6-O-, 2-O-, and 3-O-sulphotransferases) to glucosamine or uronate residues, or epimerise (C5-epimerase) β–D-glucuronate to α-L-iduronate units in the chain. Together, these enzymes produce distinct sulfation patterns both at the disaccharide level and in the completed polysaccharide. For HS, the modification enzymes act in an incomplete and interdependent fashion to form domains, consisting of regions of high sulfation flanked by intermediate sulfation ^11^. Following synthesis, the removal of 6-O-sulfate groups from the HS polysaccharide by the sulfatases (Sulf 1 and 2), may also occur ^12, 13^, potentially creating further diversity in the HS chain ^14^.

Owing to the relatively poor detection sensitivity inherent to carbohydrates compares to other biomolecules, heterogeneous HS chains are isolated from a comparatively large number of cells (typically 10^3^-10^5^ cells) or mass of tissue (typically milligrams of starting material). To advance understanding of HS structure and metabolic control mechanisms linking HS biosynthesis and expression with activity, less heterogeneous HS samples are required. This yields smaller quantities of purified material that are often beyond the limit of detection of current analysis methods ^15–21^.

Typically, the first step in HS analysis would be to obtain a disaccharide compositional profile, where disaccharides are obtained either by chemical degradation using nitrous acid, or enzymatic degradation employing bacterial lyase enzymes. The structures of the disaccharides arising from each method are distinct. The first contains intact uronate residues linked to a 2,5-anhydromannose reducing end; the second is comprised of modified (4,5-unsaturated) uronate moieties linked to an intact glucosamine reducing end, where the original identity of the uronate residue (α-L-IdoA or β-D-GlcA) has been lost. These disaccharides are termed Δ–disaccharides and have been the subject of numerous separation and labelling procedures ^16, 18, 22^, amongst which, the highest sensitivity available currently is approximately in the low femtomole (10^-15^ mol) range ^18^. Each method has its advantages and drawbacks ^23^ but all remain fundamentally limited by whatever detection system is employed.

Given that HS structure varies between cell and tissue types, even in a spatiotemporal manner, a significant advance of the sensitivity in detection of HS disaccharides is essential to enable higher resolution studies to be performed. Improved method sensitivity could conceivably translate into a detection level sufficiently low to enable the differentiation of distinct regions in individual tissues. This would complement advances in laser capture micro-dissection of tissues ^24^and cell separation and detection techniques, such as single cell analysis ^25–27^, rather than its current limitation at a relatively coarse scale.

Here, a reverse-phase ion-paired HPLC method for the separation of BODIPY-FL conjugated HS disaccharides with significantly improved detection sensitivity is presented. By employing a simple phase separation clean-up step to remove excess unreacted BODIPY-FL hydrazide and a 100-minute linear gradient, baseline separation of all 8 BODIPY-labelled HS Δ-disaccharide standards was achieved. An unprecedented practical limit of detection was achieved at less than 100 zmol (10^-21^ moles), which corresponds to ∼600 labelled molecules. The validity of the technique was confirmed first through disaccharide analysis of tinzaparin, a low molecular weight heparin of known composition ^18^, and determination of HS composition from human monocytes (demonstrating compatibility with mammalian cells). Illustration of the increased scope of HS disaccharide analysis that the improvement in sensitivity provides was then demonstrated by the investigation of HS isolated from midgut tissue of 14 *Anopheles gambiae* mosquitoes (14 midguts), a major vector for malaria in Africa. These data demonstrate unprecedented sensitivity of the method and the increased scope of HS disaccharide analysis that it affords, which is anticipated to open up many new opportunities for enhancing the toolkit for HS analysis thereby increasing understanding of HS functions in biology.

## Results and discussion

Eight major Δ–disaccharide species exist for HS (and the closely-related GAG, heparin) (Figure 1); 1-4 linked combinations of these disaccharides generate the heterogeneous nature of HS polysaccharide chains. The approach adopted here combines the use of a BODIPY-FL hydrazide fluorescent tag with reverse-phase HPLC and laser-induced fluorescence detection.

**Figure 1.**
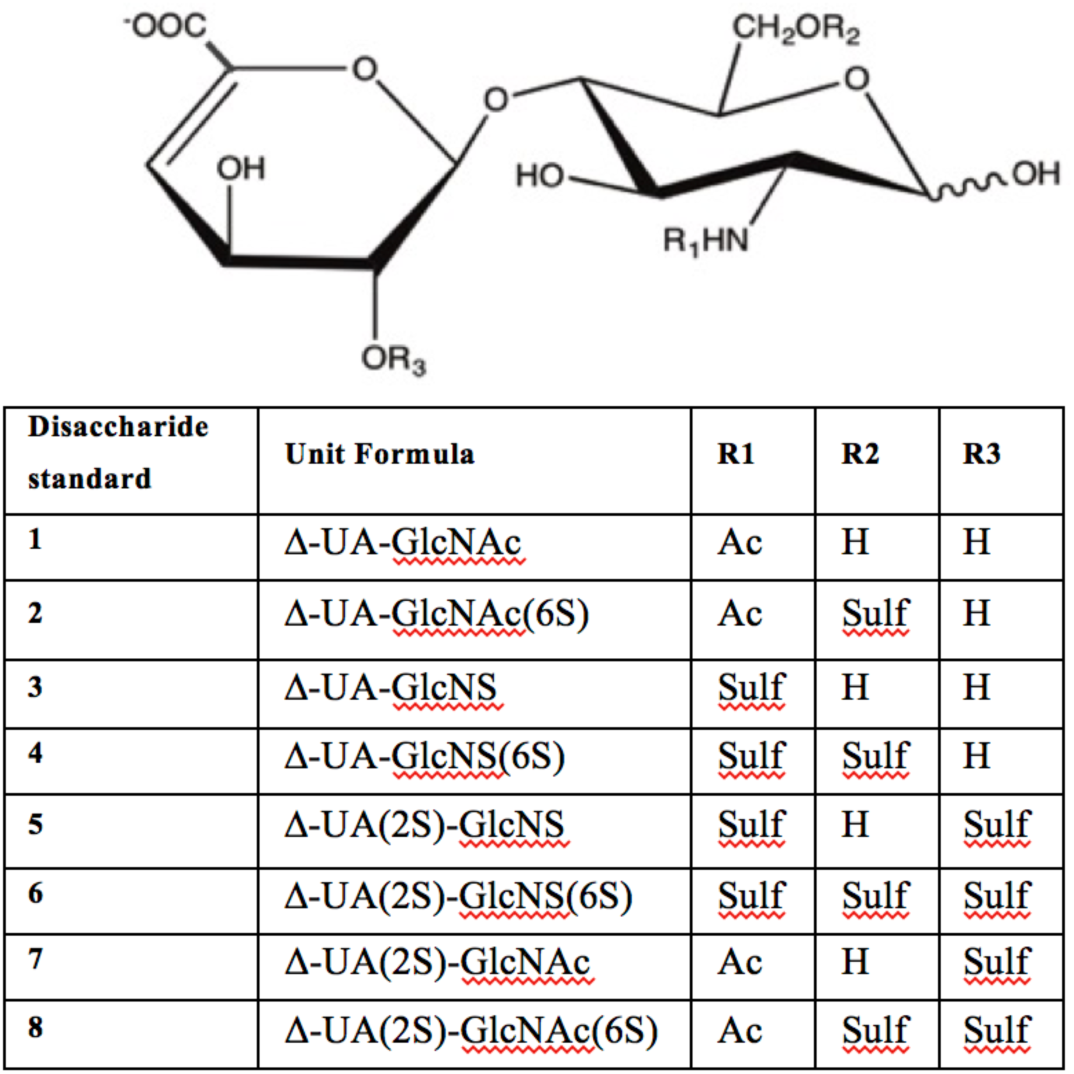
Structures of the 8 unsaturated disaccharide standards (containing ΔUA–unsaturated uronate non-reducing termini) derived from HS/heparin by exhaustive heparinase digestion (I, II and III). GlcNAc; N-acetyl-D-glucosamine, GlcNS; N-sulpho-D-glucosamine, 2S; 2-O-sulfate, 6S; 6-O-sulfate, Ac; acetyl, Sulf; sulfate, H; hydrogen.

### Removal of free BODIPY-FL hydrazide label from aqueous solution

For the highest sensitivity detection, removal of excess unreacted fluorescent tag from the labelled material without significant sample loss presented a major challenge, but was found to be essential to avoid masking of sample peaks. Here, new strategies were explored for the removal of excess BODIPY-FL hydrazide label from samples to assist separation, identification and characterisation of labelled HS disaccharides. Current methods employing BODIPY-FL rely on either thin layer chromatography (TLC), or do not attempt to remove excess fluorophore before application to the column where the majority of the BODIPY-FL hydrazide elutes at the onset of the run during the isocratic step ^15, 18^. Other fluorophores, such as 2-aminoacridone need to be pre-treated and purified before use to reduce fluorescent impurities and improve signal-to-noise ratio for detection ^16^, or may require verification, for example, by mass spectrometry ^21^. However, none of these rivals the present one in sensitivity, their best detection limit being around 10^-13^ mol. ^15, 18^. In any labelling and detection procedure, excess label remaining after the coupling reaction could co-elute with labelled species, thereby decreasing the resolving power of the method and interfering with the detection of neighbouring eluting disaccharide species. To eliminate this problem for BODIPY-FL hydrazide RP-HPLC methods, a range of organic solvents that are immiscible with water were tested for their ability to remove excess unreacted BODIPY-FL hydrazide (Figure 2A). Five of the 14 solvents used for extraction reduced aqueous fluorescence more successfully than TLC. 1,2-dichloroethane consistently and selectively removed the most fluorescence arising from the free BODIPY-FL hydrazide label (Figure 2B) and was therefore selected for application to the labelled disaccharides prior to RP-HPLC separation. Sample clean-up after BODIPY-labelling improved the baseline of the chromatogram and removed the majority of the spurious contaminating peaks, thereby enabling separation of unlabelled BODIPY-FL hydrazide from ΔUA-GlcNAc, which elutes earliest of the Δ-disaccharides in the chromatogram (Figure 2C).

**Figure 2.**
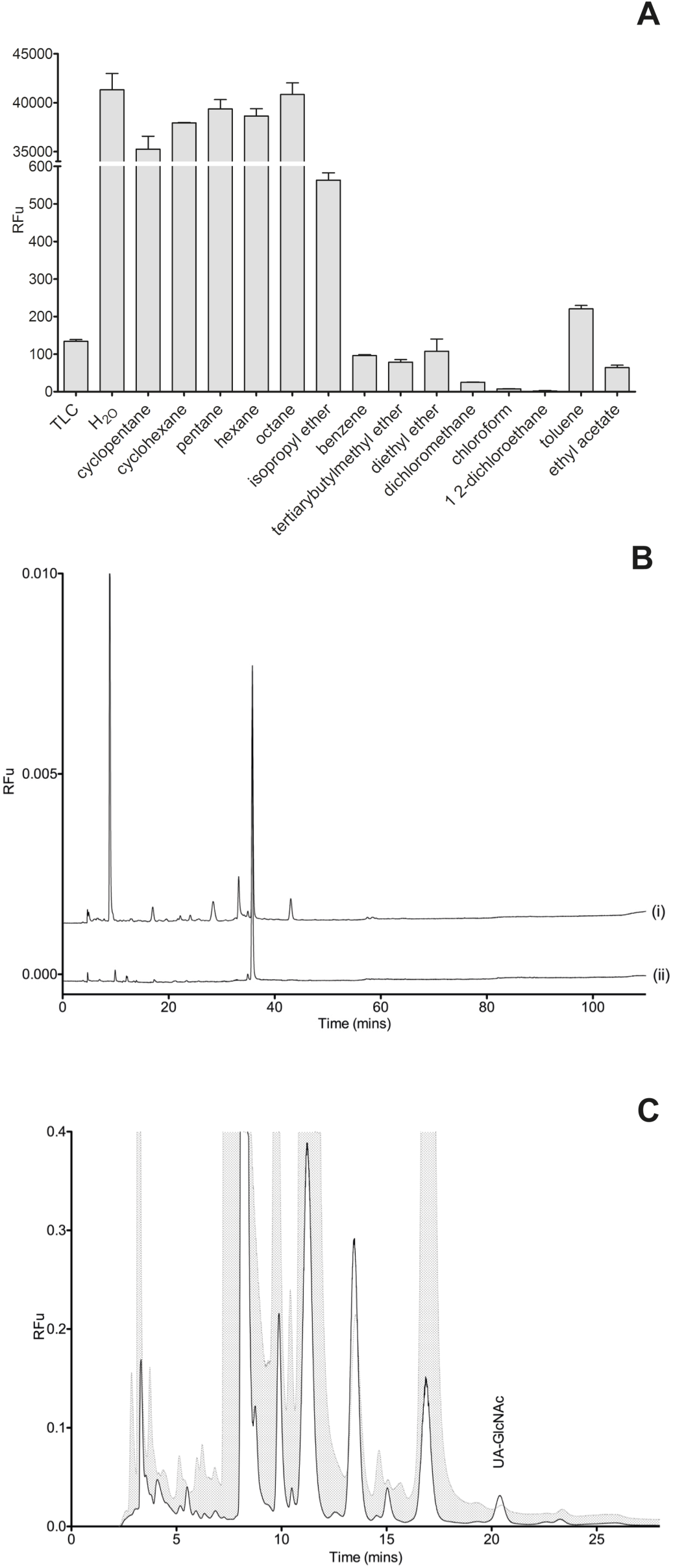
**A**. Organic solvent extraction of BODIPY-FL hydrazide label. Relative fluorescence units (RFu) of the aqueous phase following extraction of BODIPY-FL hydrazide derivatives using 14 candidate organic solvents and TLC. **B**. Ion-paired RP-HPLC of BODIPY-FL hydrazide in H_2_O after (i) TLC and (ii) 1,2-dichloroethane phase extraction. **C**. HPLC chromatogram separation of BODIPY-ΔUA-GlcNAc with and without organic solvent phase extraction. Cross hatched, without phase extraction; white in-fill, after phase extraction.

### Ion-paired RP-HPLC of HS/heparin Δ-disaccharide standards

Three commercially available 5 µm C18 silica-based columns were compared for the separation of BODIPY-labelled HS disaccharides. The separation of fluorescently-labelled material varied for each column, but exhibited similar elution profiles. The Eclipse XDB C18 column eluted fluorescent species in the shortest time period, owing to its smaller volume, but with significant peak tailing. In contrast, the SUPELCOSIL^TM^ LC18 and the ACE UltraCore Super C18 columns eluted species over a longer time period, but both exhibited augmented peak shapes for the detected eluents compared with the Eclipse XDB C18 column. The ACE UltraCore SuperC18 produced the sharpest peaks with Gaussian shapes and superior resolution of later eluting species. In addition, the ACE UltraCore SuperC18 column is stable over an extended pH range (1.5-11), facilitating method development and optimisation, as well as being compatible with liquid chromatography-mass spectrometry. Thus, the ACE UltraCore SuperC18 was selected for subsequent method optimisation. BODIPY-labelled mixtures of HS/heparin Δ-disaccharide standards were subjected to phase separation with 1,2-dichloroethane to remove excess BODIPY-FL hydrazide before resolution of all 8 disaccharides using gradient reverse phase ion-paired-HPLC (RP(IP)-HPLC) (Figures 3 & S-1).

**Figure 3.**
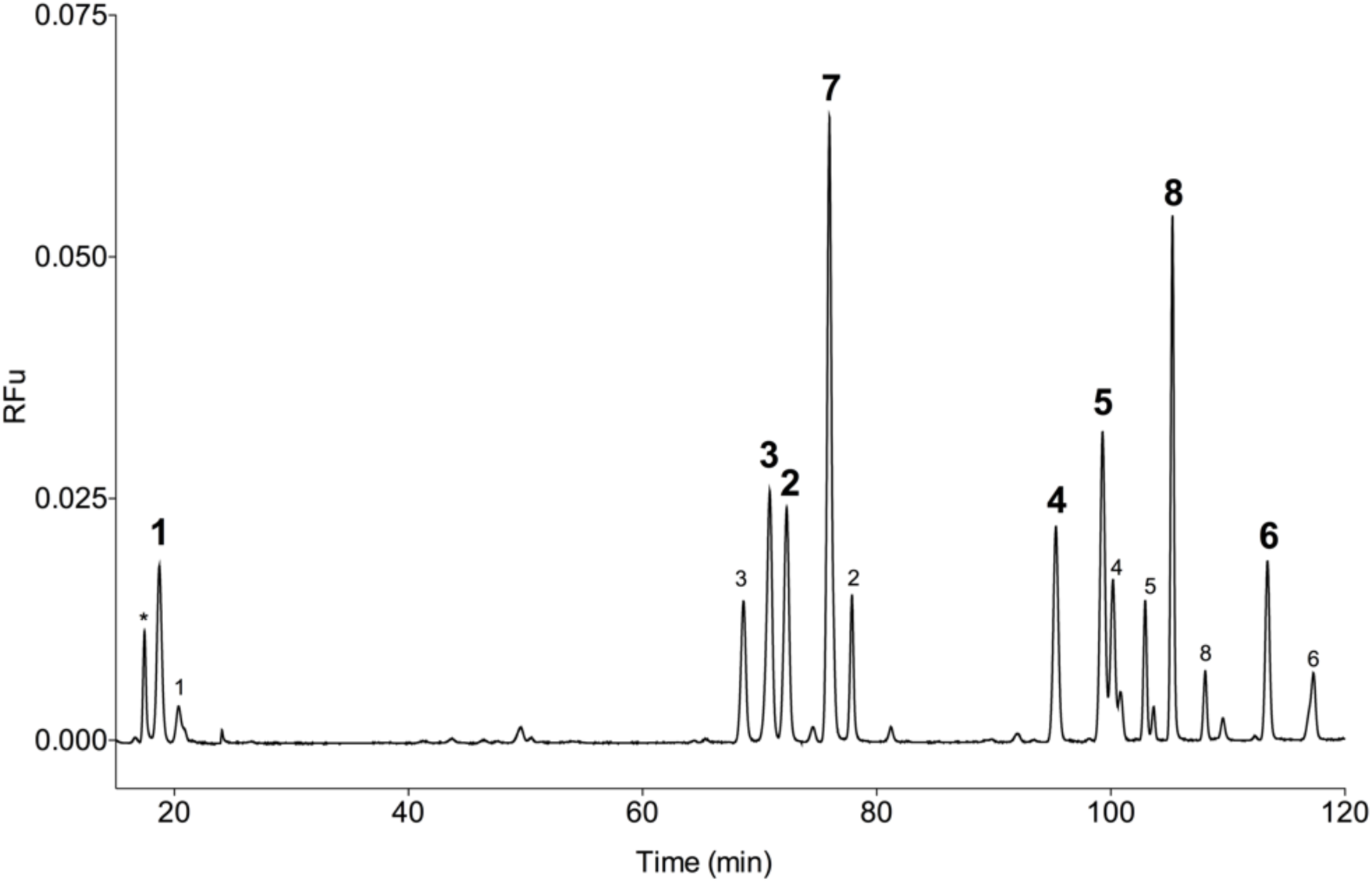
Separation of BODIPY-labelled HS/heparin Δ-disaccharide standards (1-8) by ion-paired RP-HPLC on an ACE UltraCore 5 Super C18 column (250 mm x 4.6 mm, 5 µm pore size), using a linear gradient of 10-30 mM TOAB, 0.1 M ammonium acetate, 30 % acetonitrile-100 % acetonitrile for 100 mins, following an initial 20 mins of isocratic 10 mM TOAB, 0.1 M ammonium acetate, 30 % acetonitrile. The dominant peak for each species is highlighted in bold, minor peaks from disaccharide species are also labelled. Fluorescence detection: λ_ex_ 473 nm, λ_em_ 510 nm. * unidentified peak, likely to be residual BODIPY-FL.

Several ion-pairing reagents (comprising a sequential series of tetra-butyl to heptylammonium bromide solvents, octadecyltrimethylammonium bromide and (1-dodecyl)trimethylammonium bromide) as well as methanol, and a range of pH values were also employed during method optimisation. Separation of HS/heparin disaccharides was achieved with a 100-minute gradient, using 10 mM tetraoctylammonium bromide (TOAB) in 0.1 M ammonium acetate and 30 mM TOAB in acetonitrile, delivering reproducible retention times for each peak (Table 1).

**Table 1.** Correction factors and retention times for the 8 Δ-disaccharide standards from HS/heparin. The average retention time for the dominant peak for each species and standard deviation of 4 technical replicates are shown.

| Disaccharide standard | Unit formula | Correction factor | Retention time of dominant peak (mins.secs) ±SD |
| --- | --- | --- | --- |
| 1 | ΔUA-GlcNAc | 1.00 | 21.82 ± 3.28 |
| 2 | ΔUA-GlcNAc(6S) | 0.64 | 77.19 ± 0.22 |
| 3 | ΔUA-GlcNS | 0.36 | 75.02 ± 0.57 |
| 4 | ΔUA-GlcNS(6S) | 0.20 | 94.46 ± 0.59 |
| 5 | ΔUA(2S)-GlcNS | 0.45 | 99.45 ± 0.58 |
| 6 | ΔUA(2S)-GlcNS(6S) | 0.17 | 113.19 ± 0.22 |
| 7 | ΔUA(2S)-GlcNAc | 0.27 | 80.43 ± 0.45 |
| 8 | ΔUA(2S)-GlcNAc(6S) | 1.00 | 105.00 ± 0.30 |

An inevitable consequence of the complex chemistry of the reducing end is the potential for several labelled species to be formed, resulting in complex chromatograms (Figure 3). These reactions could include at least two reaction mechanisms between sugar and label (reaction of the open-chain aldehyde with the nucleophilic fluorescent label to form a Schiff’s base or, in the case of GlcNAc residues, through reaction of the label with an oxazoline intermediate to generate an aminoglycoside ^28, 29^. Further complexity arises from the potential rearrangement of D-Glc to D-Man configuration of free reducing sugars prior to labelling following exposure to even very mild basic conditions ^30–33^. Even for seemingly simple sugars, therefore, several labelled products are usually formed and their relative proportions are difficult to predict. Consequently, the calculation and application of empirical correction factors is routinely employed in fluorophore-disaccharide methods to accommodate this variation (Table 1) ^15, 16^. The correction factor is derived from the relative labelling efficiency, which can be calculated from the average peak area of known quantities of each Δ-disaccharide standard from several runs. Where more than one peak corresponds to a Δ-disaccharide standard, the sum of the area under the peaks is used. Both the efficiency of labelling of each Δ-disaccharide standard, arising from the chemical differences in reducing end chemistry mentioned above, and the different concentration of acetonitrile that is required to elute each disaccharide contribute to the variance observed for peak values, with increased acetonitrile levels attenuating the fluorescence.

### Limit of detection

When calculating the limit of detection for a labelled substance extracted from natural sources, there are two principal considerations. The first is the amount of sample material required to permit detection, but this is a function of the particular extraction procedure used and examples of efficient extraction have been published [29]. The second consideration is the fundamental limit of dilution of the labelled material that still permits detection at an acceptable signal to noise ratio. For the present method, the practical limit of detection for all 8 Δ-disaccharide standards, by dilution from 1 nM is calculated to be less than 100 zeptomoles (100 x 10^-21^ mol) (Figure 4). This represents a dramatic improvement in the detection sensitivity of disaccharides in comparison with the fluorophore, 2-aminoacridone ^16^, and is also a marked improvement on the previous BODIPY-labelled Δ-disaccharide method (detection limit *ca*. 10^-15^ mol) ^15, 18^. In the case of the latter, the high pH (∼ pH 13) that is required to facilitate separation using HPLC-SAX partially limits the gains afforded by the high coefficient of extinction of the BODIPY-FL label when compared to other widely used fluorophores. This pH consideration precludes the use of widely available silica-based stationary phases and leads to a strong decrease in fluorescence between pH 12 and 13 (Figure S-2). The novel reverse phase methodology reported here permits the maximum sensitivity of the BODIPY-FL fluorophore to be exploited in-concert with laser-induced fluorescence, harnessing for the first time the full potential of this dye for GAG disaccharide analysis.

**Figure 4.**
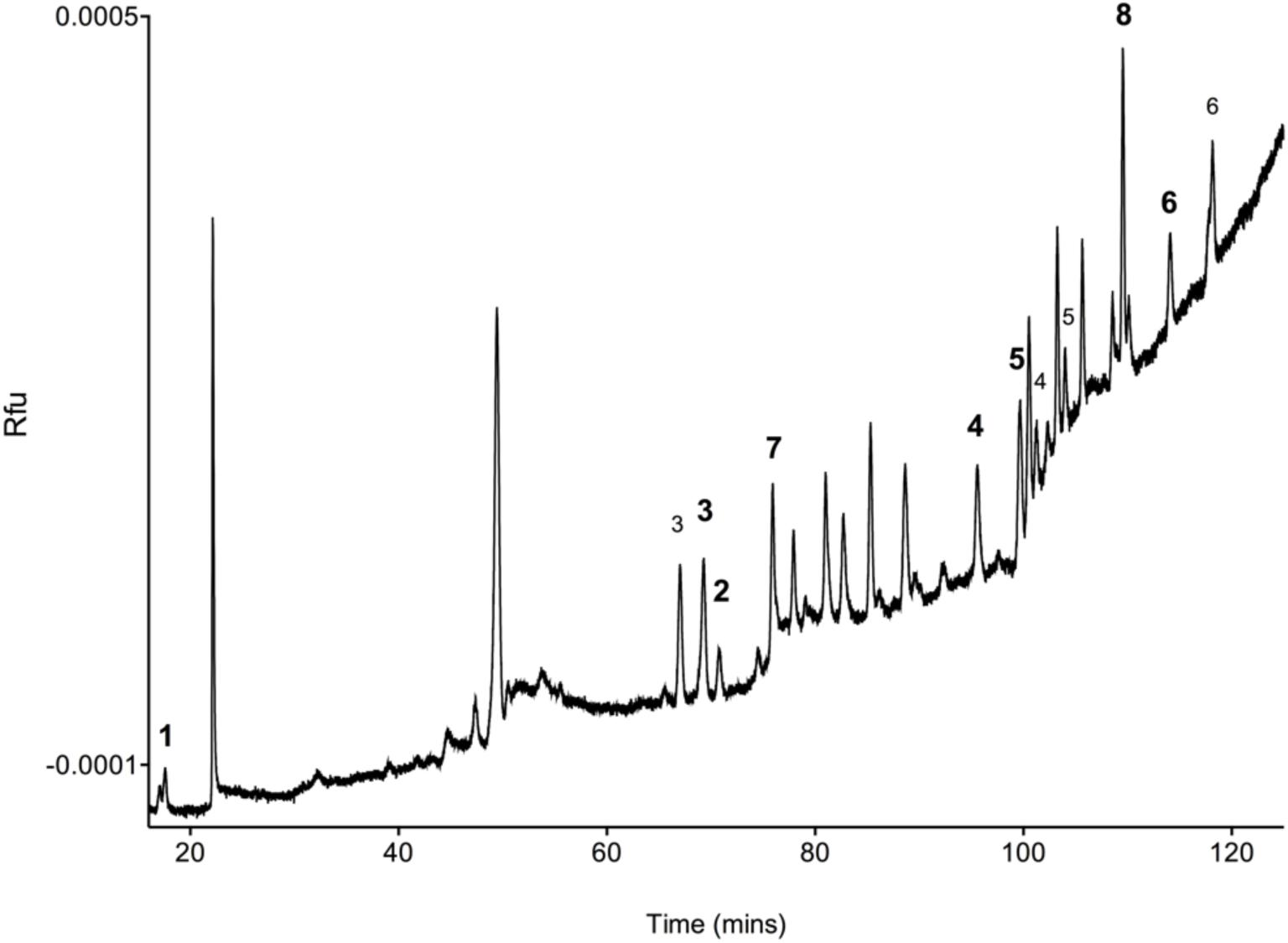
Limit of detection for all 8 HS/heparin Δ-disaccharide standards by dilution. Detection of all 8 HS/heparin disaccharides was tested by serial dilution of a labelled set of disaccharides; data are shown for 10 attogram sample. The limit of detection was around 10 attograms of loaded samples (zeptomoles, 10^-21^ moles).

### Analysis of HS/heparin from cell, tissue and commercial sources

In order to validate the method and demonstrate its utility, Δ-disaccharides derived from three different sources were analysed. Firstly, analysis of a sample of commercial tinzaparin (low molecular weight heparin) was conducted to confirm the ability of the method to yield the expected results on a sample of known composition ^18^. Tinzaparin was heparinase-digested and the disaccharide products were isolated and labelled with BODIPY-FL hydrazide. Following derivatisation with the fluorophore, separation and application of the appropriate correction factors, the predominant Δ-disaccharide species was determined to be Δ-UA(2S)-GlcNS(6S) (∼79%, Table 2), as is typical for heparins and the overall composition was consistent with the values obtained using other standard methods ^20, 34^.

**Table 2.** HS/heparin disaccharide composition analysis of tinzaparin (n=4), THP-1 monocytes (10^7^ cells extracted, n=3 biological replicates) and *A. gambiae* midguts (n=14, pooled and analysed in 2 technical replicates). nd, not detected. SEM, Standard error of the mean.

| Disaccharide species | Tinzaparin (%) ± SEM | THP-1 monocytes (%) ± SEM | *A. gambiae* midguts (%) ± SEM |
| --- | --- | --- | --- |
| ΔUA-GlcNAc | 0.66 ± 0.48 | 22.73 ± 2.65 | 17.91 ± 3.11 |
| ΔUA-GlcNAc(6S) | 0.46 ± 0.19 | 2.95 ± 0.79 | 21.90 ± 1.40 |
| ΔUA-GlcNS | 2.60 ± 0.007 | 1.56 ± 0.05 | 5.02 ± 0.40 |
| ΔUA-GlcNS(6S) | 11.71 ± 2.48 | 0.43 ± 0.43 | nd |
| ΔUA(2S)-GlcNS | 5.15 ± 1.15 | 13.03 ± 1.17 | 7.35 ± 0.39 |
| ΔUA(2S)-GlcNS(6S) | 79.19 ± 1.43 | 2.47 ± 1.07 | 22.81 ± 5.60 |
| ΔUA(2S)-GlcNAc | 0.12 ± 0.08 | 52.45 ± 5.92 | 24.56 ± 6.24 |
| ΔUA(2S)-GlcNAc(6S) | 0.11 ± 0.02 | 2.98 ± 0.10 | 0.42 ± 0.09 |
| GlcNAc | 0.66 ± 0.47 | 92.37 ± 3.30 | 64.80 ± 7.86 |
| GlcNS | 98.64 ± 0.72 | 7.63 ± 3.30 | 55.16 ± 1.30 |
| 6S | 91.46 ± 0.39 | 15.16 ± 7.47 | 45.14 ± 7.10 |
| 2S | 86.38 ± 0.02 | 65.82 ± 9.50 | 35.19 ± 7.86 |
| Un-sulfated | 0.66 ± 0.47 | 22.73 ± 3.35 | 26.29 ± 3.11 |
| Mono-sulfated | 3.18 ± 0.24 | 68.22 ± 6.78 | 63.43 ± 4.43 |
| Di-sulfated | 16.96 ± 0.72 | 6.75 ± 2.37 | 2.56 ± 1.94 |
| Tri-sulfated | 79.18 ± 1.43 | 2.30 ± 1.07 | 7.71 ± 5.61 |

Secondly, to demonstrate the compatibility of the method with a verified HS purification approach amenable for both cells and tissues ^35^, HS from human THP-1 monocytes (∼10^7^ cells) was purified and the disaccharide composition determined. As expected, human monocyte HS contained more Δ-UA-GlcNAc (∼23%) and reduced levels of Δ-UA(2S)-GlcNS(6S) (2%) than heparin, with varying levels of intermediate sulfated Δ-disaccharide species (Table 2). To the best of our knowledge, this represents the first report of HS Δ-disaccharide composition for THP-1 cells, a monocytic cancer cell line widely used for immunological studies *in vitro*. HS is the major GAG in THP-1 cell membranes ^36^ with a distinct profile compared to peripheral blood mononuclear cells ^19, 36^. The major disaccharide species was determined to be Δ-UA(2S)-GlcNAc (∼52%). This is unusually high compared to other types of cell and tissue HS composition. Since Δ-UA(2S)-GlcNAc disaccharide percentages for tinzaparin (∼0.12%) and *A. gambiae* midgut (∼25%) HS using the same method did not mirror the THP-1 results, it is unlikely that bias in method analysis is responsible for the high percentage of this particular (usually relatively rare) disaccharide. Moreover, the THP-1 HS was prepared using the same approach as the midgut tissue and other HS profiles published previously ^35, 37–39^. Thus conceivably, the high prevalence of Δ-UA(2S)-GlcNAc could be a specific feature of THP-1 cellular HS, attributed to its adaptation to cell culture conditions or cancerous origin.

Thirdly, HS was isolated from an *in vivo* tissue source (mosquito midgut tissue; 14 samples, 850 µg (wet weight) starting material), using pre-established methods ^15, 35^ and subjected to disaccharide analysis after digestion with multiple heparinases as described earlier. Peak detection was achieved comfortably above the limit of detection for seven of the 8 HS disaccharide species, suggesting that analysis should be possible from individual midguts. This would enable population diversity of HS within individual mosquitoes midguts to be examined. The percentage contribution of each HS Δ-disaccharide species was calculated as for the other HS/heparin samples. Mosquito midgut HS contained ∼18% Δ-UA-GlcNAc and ∼23% Δ-UA(2S)-GlcNS(6S) (Table 2). The mono-sulfated species, Δ-UA(2S)-GlcNAc (∼25%) and Δ-UA-GlcNAc(6S) (∼22%) were also prominent suggesting a different compositional HS domain structure than either heparin or THP-1 monocyte HS, although the percentage contribution of mono-sulfated disaccharides overall for THP-1 and midgut HS were similar (∼68%, and ∼63% respectively). The mosquito species, *A. gambiae*, is the main vector for malaria in Africa. Malaria parasites invade the mosquito midgut wall where they transform from ookinetes into sporozoites that then migrate to the salivary glands and are injected into humans during a bloodmeal. A tissue specific HS profile for mosquito midguts has been reported for the major malarial vector in India, *A. stephensi* ^17^. Interestingly, data presented here for *A. gambiae* suggest that *A. gambiae* midgut HS is distinct from that reported for *A. stephensi*, indicating that HS composition may vary between malaria vector species. Notably, 247 mosquito midguts (3.7 mg (dry mass) starting material) were required for the HS analysis of *A. gambiae* performed by Sinnis *et. al.*, compared to just 14 midguts (850 µg (wet weight) starting material) utilised for the results reported herein. Furthermore, for *A. stephensi*, the method detected only 6 of the 8 common HS disaccharides in human, suggesting the other two may either be below the limit of detection or are not present in *A. stephensi* midgut HS. The method reported here detected 7 of the 8 disaccharides. The increased sensitivity demonstrated by this method afforded the detection of disaccharides often reported to be low in abundance such as Δ-UA(2S)-GlcNAc(6S) in tinzaparin 0.1% and mosquito midgut (∼0.4%) and Δ-UA(2S)-GlcNAc in tinzaparin (0.1%), which is not always possible by other established methods ^17^. Therefore, this method will be invaluable in the near future to detect other rare HS disaccharide species (i.e. 3-O-sulfated disaccharides) once authentic disaccharide standards become commercially available for their analysis.

The separation and improved detection sensitivity of this method will facilitate the development of sequencing and structural analyses for HS and its close relative, heparin, as well as other GAGs. The reverse phase separation conditions are also compatible with mass spectrometry, as well as nano- and micro-HPLC methodologies.

## Conclusions

An ultra-high sensitivity RP(IP)-HPLC method has been developed for the separation of BODIPY-labelled HS/heparin disaccharides providing significant (from *ca.* 10^-15^ to 10^-21^ mol) sensitivity enhancement over existing techniques. The RP(IP)-HPLC method enables high sensitivity detection using standard binary HPLC equipment combined with commercially available LIF detector and standard reverse phase columns. The sensitivity achievable was demonstrated through effective HS disaccharide analysis from small amounts of *in vivo* tissue from mosquito midguts, without approaching the limit of detection.

The present method also avoids the use of high pH, which is known to reduce fluorescence intensity, require expensive polymer-based SAX chromatography and can introduce modifications to the structure of GAGs ^30–32^ that lead to further spurious peaks in the chromatograms. This method is also compatible with base-sensitive chemical derivatives such as benzoyl esters, which are employed during the production of some commercial pharmaceutical heparin samples. All 8 major Δ-disaccharide standards from heparin and HS are resolved using a 100-minute gradient with a simple phase extraction step prior to separation, with significantly improved sensitivity for the detection of small quantities of HS and heparin Δ-disaccharide material. This significantly improved sensitivity enables small amounts of cultured cell- and tissue-extracted HS/heparin samples to be analysed, greatly increasing the scope of HS structural analysis and opening up new potential experimental avenues. Furthermore, the use of volatile solvent and NaCl-free conditions facilitates BODIPY-labelled disaccharide technology for online mass spectrometry, as well as permitting adaptation to nano- and micro-HPLC systems. This could be envisaged to support future development of advanced methods for analysis and sequencing of HS and other GAG oligosaccharides and the detailed exploration of structure-activity relationships.

## Methods

### General

All reagents were purchased from Sigma-Aldrich unless specified.

### Organic solvent and thin layer chromatography extraction of excess BODIPY-FL hydrazide

1 µl BODIPY-FL hydrazide (5 mg/mL, Setareh Biotech, Eugene, OR, USA) in DMSO was diluted in HPLC grade water before addition of organic solvent in a 1:9 (v/v) ratio, followed by brief vortexing and recovery of the aqueous phase (repeated 5 times). Thin layer chromatography (TLC) was subsequently performed, in which 1 µl of BODIPY-FL hydrazide (5 mg/mL) was spotted onto foil backed TLC silica, air-dried and then developed in butan-1-ol as the mobile phase (5 ascents with drying between each ascent). The silica media was dislodged from the foil, suspended in 1 mL H_2_O and filtered using a 0.2 µm centrifugal filter to recover the sample. 200 µl of the filtrate was analysed in a black 96-well plate (Corning) for fluorescence (λ_ex_ 488 nm, λ_em_ 520 nm) and 40 µl was separated by ion-paired RP-HPLC using an ACE UltraCore 5 SuperC18 column (250 mm x 4.6 mm, 5 μm, Hichrom) equilibrated in solvent A (0.1 M ammonium acetate, 10 mM tetraoctylammonium bromide (TOAB), 30% acetonitrile) with a linear gradient of 0 - 100% solvent B (acetonitrile, 30 mM TOAB) over 120 mins.

### BODIPY-FL fluorescence in different pH conditions

HPLC grade water was adjusted incrementally through the pH range 3-6 using HCl and pH 8-13 using NaOH. HPLC grade acetonitrile was serially diluted 1:2 (v/v) with HPLC grade water. 1 µL BODIPY-FL hydrazide (5 µg/µL in DMSO) was added in triplicate to 100 µL of each condition in a black 96-well plate. Fluorescence (λex 488 nm, λem 520 nm) was measured using an Infinite M200 Pro (Tecan) instrument during experiments examining the effects of pH and with scanning (λ_ex_ 488 nm, λ_em_ 502-550 nm) for investigation of the effects of acetonitrile/water conditions.

### BODIPY-labelling of disaccharides at the reducing end

Δ-disaccharides (>95% purity, Iduron, Alderley Edge, UK) from heparin/HS (Figure 1) were labelled with BODIPY-FL hydrazide as previously described ^40^ with the omission of the reducing step. Briefly, lyophilised disaccharides were re-suspended in 10 µL BODIPY-FL hydrazide (5 mg/mL) in 85% DMSO / 15% ethanoic acid (v/v) at room temperature for 4 hours. Labelled samples were then lyophilised and re-suspended in 100 µL of HPLC grade water before phase extraction of excess BODIPY-FL hydrazide label using sequential extraction with 1,2-dichloroethane (5 x 1 mL) in a glass tube.

### Ion-paired RP-HPLC separation of Δ-disaccharides

BODIPY-labelled Δ-disaccharides were resolved using a standard binary HPLC system (Cecil, Cambridge, UK) equipped with either an ACE UltraCore 5 SuperC18 column (250 mm x 4.6 mm, 5 μm, Hichrom), SUPELCOSIL LC18 (30 cm x 4 mm, 5 μm, Sigma-Aldrich), or an Eclipse XDB-C18 column (150 mm x 4.6 mm, 5 μm Agilent technologies) and an in-line fluorescence detector (λ_ex_ 473 nm, λ_em_ 510 nm, Picometrics, Toulouse, France) under the following conditions: isocratic 100% A at a flow rate of 0.5 mL/min for 20 mins, then linear gradient elution of 0-100% B at a flow rate of 0.5 mL/min for 100 mins, where solvent A contained 0.1 M ammonium acetate, 30% HPLC grade acetonitrile, 10 mM tetraoctylammonium bromide (TOAB) and solvent B contained 30 mM TOAB dissolved in HPLC grade acetonitrile (VWR). The column was subsequently cleaned for 10 mins using solvent B (isocratic) before re-equilibration with solvent A for 10 mins between separations.

### Preparation of tinzaparin Δ-disaccharides

Tinzaparin (5 mg, EDQM (Conseil de l’Europe)) was lyophilised and digested with a cocktail of heparinases (I, II, III) (Iduron, Alderley Edge, UK) in 100 mM sodium acetate, 10 mM calcium acetate, pH 7 for 24 hours at 37℃. Post digestion, the samples were incubated at 95 ℃ for 5 mins to ablate enzyme function. The digest was applied to a column (1000 mm x 300 mm) of BioGel P6 resin (Bio-Rad, UK) for size exclusion chromatography in isocratic 0.25 M ammonium chloride (Fisher, UK) at a flow rate of 0.2 mL/min. Elution of the Δ-disaccharide material from the column was monitored by absorbance of the 4,5 carbon double bond (λ_abs_ = 232 nm) introduced by the heparinase enzyme digestion. The disaccharide fraction was collected and desalted using a Sephadex-G10 column (GE Healthcare Life Sciences) at a flow rate of 2 mL/min in HPLC grade water before lyophilisation and BODIPY-FL hydrazide labelling shown above.

### Preparation of THP-1 monocyte HS Δ-disaccharides

THP-1 monocytes were cultured in RPMI-1640 (Gibco) supplemented with 10% foetal bovine serum and 2 mM L-glutamine at 37℃ in 5% CO_2_ conditions. Cells were washed with PBS and re-suspended in 1% Triton X-100/PBS to solubilise HS proteoglycans. Proteins were digested with 2mg/mL Pronase in 100 mM tris acetate, 10 mM calcium acetate buffer pH 5 for 4 hours at 37℃. Anion exchange chromatography was performed using DEAE beads (Sigma) as previously described 25 and eluted samples were desalted using PD10 columns according to the manufacturer’s instructions (GE Healthcare). Samples were applied to centrifuge filters (Vivaspin, MWCO 5,000) and washed with HPLC grade water. The retentate was lyophilised before HS enzyme digestion with heparinases I, II and III in 0.1 M sodium acetate, 0.1 mM calcium acetate, pH 7.0 for 16 hours at 37°C. Digestions were lyophilised and labelled with BODIPY-FL hydrazide.

### Preparation of Anopheles gambiae midgut HS Δ-disaccharides

The *A. gambiae* colony used in this study was the G3 strain, originally established from mosquitoes collected in Gambia and maintained under laboratory conditions for several decades. Adult mosquitoes were maintained in small BugDorm cages (17.5 x 17.5 x 17.5 cm) in the insectary at Keele University, under a 12/12 hour light/dark photoperiod at 27°C with 75% humidity. Larvae were fed on TetraMin tropical fish food flakes (Tetra) and adult mosquitoes were allowed to feed *ad libitum* on a 10% (w/v) sugar solution. Midguts from female *A. gambiae* were dissected and placed in 0.5 mL TRIzol (Thermo fisher) on ice. HS extraction and purification was conducted as previously described ^35^ and the resultant HS disaccharides were labelled with BODIPY-FL hydrazide.

## Acknowledgements

This work was funded by BBSRC and EPSRC grants no. BB/L023717/1 (MAS & MLMH), BB/M019209/1 (MAS), the Royal Society of Tropical Medicine and Hygiene Small Grant 2016-17 (MLMH), and a Biochemical Society Vacation studentship (AL and AKP).

## Author contributions

MAS, AKP, EAY, JET and MLMH designed the approach and interpreted the results. MAS, AKP, EAY, JET, SEG, ACNL, AL and MLMH defined the method, and MLMH implemented it. MAS, AKP, EAY and MLMH analysed the data. MAS and AKP supervised the study. All of the authors drafted and approved the manuscript. The manuscript was written by MAS, EAY, JET and MLMH. All authors have given approval to the final version of the manuscript.

## Supplementary Figures

**Figure S-1.**
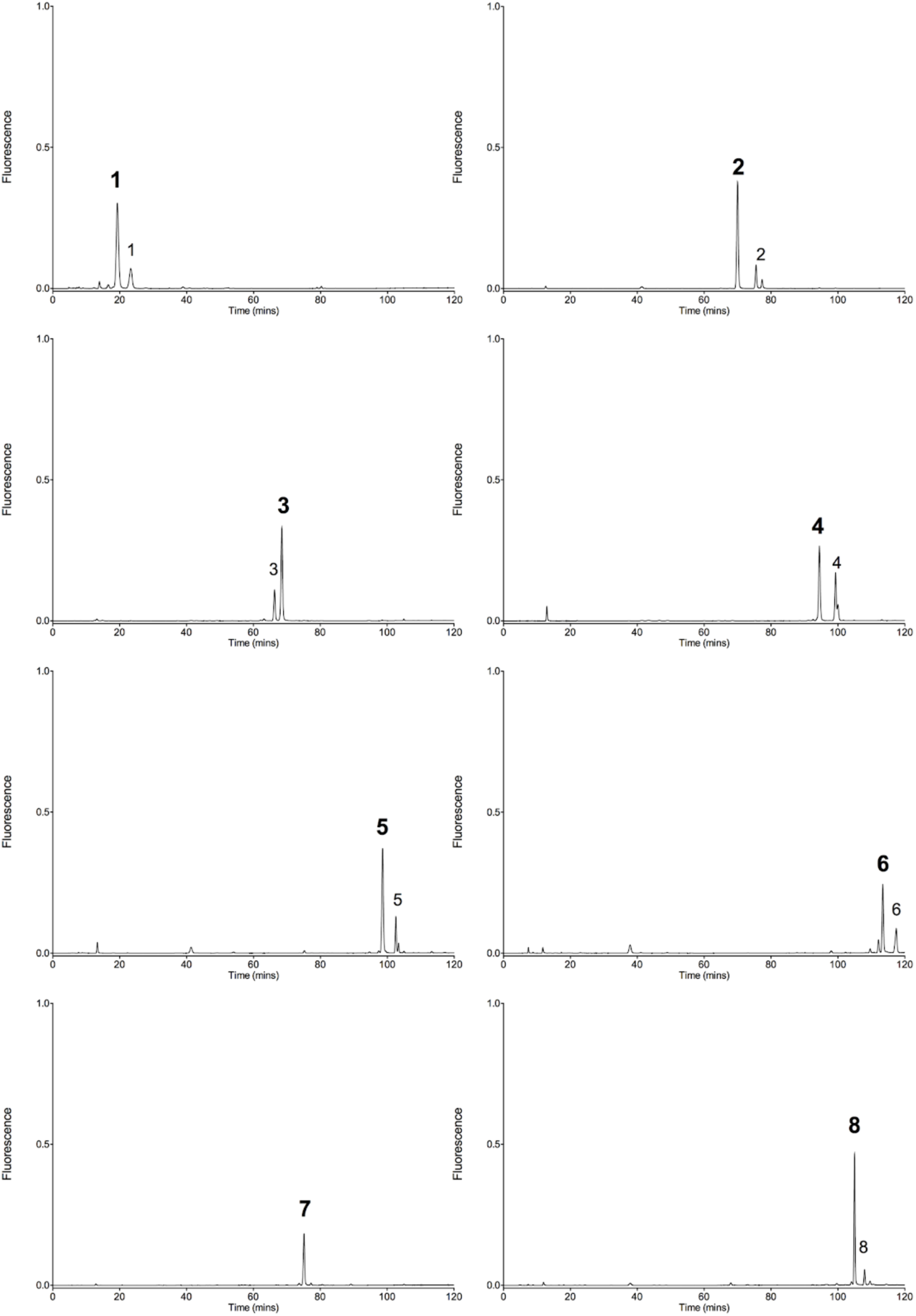
Chromatograms of BODIPY-labelled individual HS disaccharides 1-8. Dominant peaks are noted in bold, minor peak species are also labelled.

**Figure S-2.**
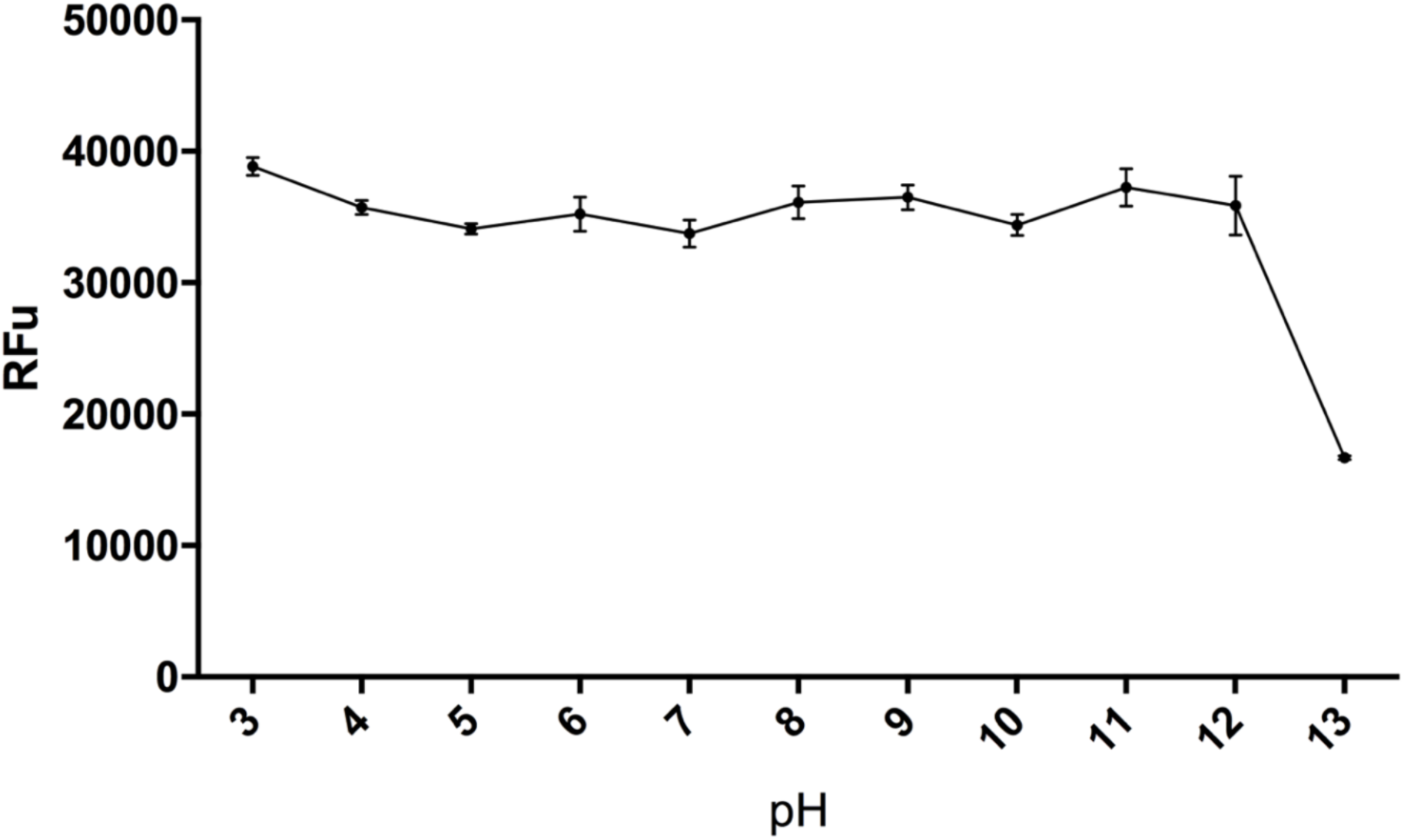
BODIPY-FL hydrazide fluorescence in water pH 3-13. Error bars represent the SEM of three replicates.

## References

1. Connell, B.J. & Lortat-Jacob, H. Human immunodeficiency virus and heparan sulfate: from attachment to entry inhibition. Frontiers in Immunology 4(2013).

2. Kim SY, et al. Interaction of Zika Virus Envelope Protein with Glycosaminoglycans. Journal of Biochemistry, DOI: 10.1021/acs.biochem.1026b01056. (2017).

3. Ghezzi S, et al. Heparin prevents Zika virus induced-cytopathic effects in human neural progenitor cells. Journal of Antiviral Research 140, 13–17 (2017).

4. Carruthers, V.B., Hakansson, S., Giddings, O.K. & Sibley, L.D. Toxoplasma gondii uses sulfated proteoglycans for substrate and host cell attachment. Infection and Immunity 68, 4005–4011 (2000).

5. Zhang, Y., et al. A comparative study on the heparin-binding proteomes of Toxoplasma gondii and Plasmodium falciparum. Proteomics 14, 1737–1745 (2014).

6. Frevert, U., et al. MALARIA CIRCUMSPOROZOITE PROTEIN BINDS TO HEPARAN-SULFATE PROTEOGLYCANS ASSOCIATED WITH THE SURFACE-MEMBRANE OF HEPATOCYTES. Journal of Experimental Medicine 177, 1287–1298 (1993).

7. Butcher, B.A., Sklar, L.A., Seamer, L.C. & Glew, R.H. HEPARIN ENHANCES THE INTERACTION OF INFECTIVE LEISHMANIA-DONOVANI PROMASTIGOTES WITH MOUSE PERITONEAL-MACROPHAGES - A FLUORESCENCE FLOW CYTOMETRIC ANALYSIS. J. Immunol. 148, 2879–2886 (1992).

8. Fatoux-Ardore, M., et al. Large-Scale Investigation of Leishmania Interaction Networks with Host Extracellular Matrix by Surface Plasmon Resonance Imaging. Infection and Immunity 82, 594–606 (2014).

9. Maciej-Hulme, M.L., Skidmore, M.A. & Price, H.P. The role of heparan sulfate in host macrophage infection by Leishmania species. Biochem Soc Trans 46, 789–796 (2018).

10. Goode, K.J., Poon, I.K.H., Phipps, S. & Hulett, M.D. Soluble Heparan Sulfate Fragments Generated by Heparanase Trigger the Release of Pro-Inflammatory Cytokines through TLR-4. Plos One 9(2014).

11. Turnbull, J.E. & Gallagher, J.T. DISTRIBUTION OF IDURONATE 2-SULFATE RESIDUES IN HEPARAN-SULFATE - EVIDENCE FOR AN ORDERED POLYMERIC STRUCTURE. Biochem. J. 273, 553–559 (1991).

12. Lamanna, W.C., et al. Heparan sulfate 6-O-endosulfatases: discrete in vivo activities and functional co-operativity. Biochemical Journal 400, 63–73 (2006).

13. Frese, M.A., Milz, F., Dick, M., Lamanna, W.C. & Dierks, T. Characterization of the Human Sulfatase Sulf1 and Its High Affinity Heparin/Heparan Sulfate Interaction Domain. Journal of Biological Chemistry 284, 28033–28044 (2009).

14. Yates, E.A., Gallagher, J.T. & Guerrini, M. Introduction to the Molecules Special Edition Entitled’. Molecules 24(2019).

15. Skidmore, M.A., et al. High sensitivity separation and detection of heparan sulfate disaccharides. Journal of Chromatography A 1135, 52–56 (2006).

16. Deakin, J.A. & Lyon, M. A simplified and sensitive fluorescent method for disaccharide analysis of both heparan sulfate and chondroitin/dermatan sulfates from biological samples. Glycobiology 18, 483–491 (2008).

17. Sinnis, P., et al. Mosquito heparan sulfate and its potential role in malaria infection and transmission. Journal of Biological Chemistry 282, 25376–25384 (2007).

18. Skidmore, M.A., Guimond, S.E., Dumax-Vorzet, A.F., Yates, E.A. & Turnbull, J.E. Disaccharide compositional analysis of heparan sulfate and heparin polysaccharides using UV or high-sensitivity fluorescence (BODIPY) detection. Nature Protocols 5, 1983–1992 (2010).

19. Shao, C., et al. Comparative glycomics of leukocyte glycosaminoglycans. Febs J 280, 2447–2461 (2013).

20. Galeotti, F. & Volpi, N. Novel reverse-phase ion pair-high performance liquid chromatography separation of heparin, heparan sulfate and low molecular weight-heparins disaccharides and oligosaccharides. Journal of Chromatography A 1284, 141–147 (2013).

21. Volpi, N., Galeotti, F., Yang, B. & Linhardt, R.J. Analysis of glycosaminoglycan-derived, precolumn, 2-aminoacridone-labeled disaccharides with LC-fluorescence and LC-MS detection. Nature Protocols 9, 541–558 (2014).

22. Galeotti, F. & Volpi, N. Online Reverse Phase-High-Performance Liquid Chromatography-Fluorescence Detection-Electrospray Ionization-Mass Spectrometry Separation and Characterization of Heparan Sulfate, Heparin, and Low-Molecular Weight-Heparin Disaccharides Derivatized with 2-Aminoacridone. Analytical Chemistry 83, 6770–6777 (2011).

23. Powell, A.K., Yates, E.A., Fernig, D.G. & Turnbull, J.E. Interactions of heparin/heparan sulfate with proteins: Appraisal of structural factors and epxerimental approaches. Glycobiology 14, 17R–30R (2004).

24. Nagai-Okatani, C., Nagai, M., Sato, T. & Kuno, A. An Improved Method for Cell Type-Selective Glycomic Analysis of Tissue Sections Assisted by Fluorescence Laser Microdissection. Int J Mol Sci 20(2019).

25. La Manno, G., et al. Molecular Diversity of Midbrain Development in Mouse, Human, and Stem Cells. Cell 167, 566–580.e519 (2016).

26. Dou, M., et al. High-Throughput Single Cell Proteomics Enabled by Multiplex Isobaric Labeling in a Nanodroplet Sample Preparation Platform. Anal Chem 91, 13119–13127 (2019).

27. Zhu, Y., et al. Single-cell proteomics reveals changes in expression during hair-cell development. Elife 8(2019).

28. Jha, R. & Davis, J.T. Hydrolysis of the GlcNAc oxazoline: deamidation and acyl rearrangement. Carbohydr Res 277, 125–134 (1995).

29. Makino, A. & Kobayashi, S. Chemistry of 2-oxazolines: A crossing of cationic ring-opening polymerization and enzymatic ring-opening polyaddition. Vol. 48 1251-1170 (J poly Sci(A), 2010).

30. Angyal, S. The Lobry de Bruyn-Alberda van Ekenstein transformation and related reactions, in: Glycoscience: epimerisation, isomerisation and rearrangement reactions of carbohydrates. Vol. 215 1–14 (Springer-Verlag Berlin, 2001).

31. Lobry de Bruyn, C. & van Ekenstein, W. Action of alkalis on the sugars. Reciprocal transformation of glucose, fructose and mannose., Vol. 14 203–216 (Rec.Trav.Chim. Pays-Bas.,, 1895).

32. Yamada, S., Watanabe, M. & Sugahara, K. Conversion of N-sulfated glucosamine to N-sulfated mannosamine in an unsaturated heparin disaccharide by non-enzymatic, base-catalyzed C-2 epimerization during enzymatic oligosaccharide preparation. Carbohydrate Research 309, 261–268 (1998).

33. Toida, T., Vlahov, I.R., Smith, A.E., Hileman, R.E. & Linhardt, R.J. C-2 epimerization of N-acetylglucosamine in an oligosaccharide derived from heparan sulfate. J Carbohyd Chem 15, 351–360 (1996).

34. Yang, B., et al. Ultra-performance ion-pairing liquid chromatography with on-line electrospray ion trap mass spectrometry for heparin disaccharide analysis. Anal Biochem 415, 59–66 (2011).

35. Guimond, S.E., et al. Rapid Purification and High Sensitivity Analysis of Heparan Sulfate from Cells and Tissues TOWARD GLYCOMICS PROFILING. Journal of Biological Chemistry 284, 25714–25722 (2009).

36. Makatsori, E., et al. Synthesis and distribution of glycosaminoglycans in human leukemic B- and T-cells and monocytes studied using specific enzymic treatments and high-performance liquid chromatography. Biomed Chromatogr 15, 413–417 (2001).

37. Chan, W.K., et al. 2-O Heparan Sulfate Sulfation by Hs2st Is Required for Erk/Mapk Signalling Activation at the Mid-Gestational Mouse Telencephalic Midline. PLoS One 10, e0130147 (2015).

38. Kalus, I., et al. Sulf1 and Sulf2 Differentially Modulate Heparan Sulfate Proteoglycan Sulfation during Postnatal Cerebellum Development: Evidence for Neuroprotective and Neurite Outgrowth Promoting Functions. PLoS One 10, e0139853 (2015).

39. O’Neill, P., et al. Sulfatase-mediated manipulation of the astrocyte-Schwann cell interface. Glia 65, 19–33 (2017).

40. Skidmore, M.A., et al. High sensitivity separation and detection of heparan sulfate disaccharides. J Chromatogr A 1135, 52–56 (2006).

